# Plan versus motion-referenced generalization of fast and slow processes in reach adaptation

**DOI:** 10.1101/2022.07.13.499894

**Authors:** Judith L. Rudolph, Luc P.J. Selen, W. Pieter Medendorp

**Affiliations:** Radboud University, Donders Institute for Brain, Cognition and Behaviour, Nijmegen, The Netherlands

**Keywords:** Motor learning, reaching, state-space modeling, motor primitives, generalization

## Abstract

Generalization in motor learning refers to the transfer of a learned compensation to other relevant contexts. The generalization function is typically assumed to be of Gaussian shape, centered on the planned motion, although more recent studies associate generalization with the actual motion. Because motor learning is thought to involve multiple adaptive processes with different time constants, we hypothesized that these processes have different time-dependent contributions to the generalization. Guided by a model-based approach, the objective of the present study was to experimentally examine these contributions. We first reformulated a validated two-state adaptation model as a combination of weighted motor primitives, each specified as a Gaussian-shaped tuning function. Adaptation in this model is achieved by updating individual weights of the primitives of the fast and slow adaptive process separately. Depending on whether updating occurred in a plan-referenced or a motion-referenced manner, the model predicted distinct contributions to the overall generalization by the slow and fast process. We tested 23 participants in a reach adaptation task, using a spontaneous recovery paradigm consisting of five successive blocks of a long adaptation phase to a viscous force field, a short adaptation phase with the opposite force, and an error-clamp phase. Generalization was assessed in eleven movement directions relative to the trained target direction. Results of our participant population fell within a continuum of evidence for plan-referenced to evidence for motion-referenced updating. This mixture may reflect the differential weighting of explicit and implicit compensation strategies among participants.

## Introduction

We continuously adapt our movements to accommodate for varying dynamics of the body and the environment. During this adaptation, our brain adjusts its internal model of these dynamics based on observed errors and uses this model in future transformations between motor commands and sensory outcomes. The updated internal model not only accounts for the situation experienced, but also affects sensorimotor performance in similar situations, called generalization.

It has long been thought that internal models are updated based on a combination of the movement errors experienced and the movement plans (as defined by the goal) that led to those errors (Donchin et al., 2003; Thoroughman & Shadmehr, 2000; Thoroughman & Taylor, 2005), referred to as plan-referenced learning. However, more recent work by Gonzalez Castro et al. (2011) has shown that the patterns of generalization associated with alternating force field adaptation of reaching movements are associated with the actual, not the planned motion, with maximal generalization where actual motions were clustered due to the perturbations. This suggests that the combination of the errors experienced and the experienced motion states (defined by the effector or muscles), rather than the planned motion states, drives the adaptation, referred to as motion-referenced learning.

It has also been argued that the error-based updating of the internal model is not simply governed by a single adaptive process, but involves multiple processes with different time constants. Fast processes learn quickly from errors but also forget rapidly, whereas slow processes learn slowly and hardly forget (Inoue et al., 2015; Lee & Schweighofer, 2009; Smith et al., 2006). Signatures of these different time scales can be revealed using spontaneous recovery paradigms: after long exposure to one type of perturbation and a much shorter exposure to the opposite perturbation, participants show a rebound to the adaptation for the initial perturbation. Smith et al. (2006) explained this observation by a two-state adaptation model, in which the brief reverse-adaptation is driven by a fast adaptive state and the rebound mainly follows from the lagging slow adaptive state, as built up during the initial adaptation and retained during the reverse-adaptation. These two processes, which together contribute to the update of the internal model, do not only explain spontaneous recovery (Ethier et al., 2008), but also behavioral observations such as faster re-relearning/re-adaptation (Kojima et al., 2004) and interference (Davidson & Wolpert, 2004).

In neurocomputational terms, the implementation of the internal model is conceived as a network of motor primitives, each specified as a Gaussian-like tuning function (Thoroughman & Shadmehr, 2000; Yokoi et al. 2011, Yokoi et al. 2014). These primitives are weighted and combined to generate the control signals for the task goal. Adaptation is achieved by error-based updating of these weights. However, how the fast and slow adaptive processes contribute to this updating is not known, let alone whether they operate by crediting the planned or actual motion. The objective of the present study is to investigate the contribution of the slow and fast process to the overall generalization in a spontaneous recovery paradigm using a force field adaptation task.

Modeling work by Tanaka and colleagues suggests that the fast process, due to its strong sensitivity to single trial error, contributes mostly to trial-by-trial generalization, whereas the slow process, due to its strong retention, contributes more to post-adaptation generalization (Tanaka et al., 2012). Yet, these authors did not consider that adaptive processes may be driven by different credit assignment of errors. For example, studies on visuomotor adaptation have dissociated the fast and slow adaptative process in terms of explicit versus implicit learning (Day et al., 2016; McDougle et al., 2015, 2017), which contributes to the notion that they attribute errors to different aspects of the motion.

Here, we modified the dual-rate model by Smith et al. (2006) to account for generalization, by separately updating motor primitive weights for the fast and the slow process (Donchin et al., 2003; Thoroughman & Shadmehr, 2000). The location of this weight updating was either centered on the planned motor states (plan-referenced) or on the actually experienced motor states (motion-referenced), leading to distinct contributions of the slow and fast process to the overall generalization pattern.

If both adaptive processes credit the planned movement, during the exposure to the first force field their generalization contributions build up around the target location, with different time constants (see Fig 1B, upper row). The overall generalization is the sum of these two contributions. During exposure to the second force field, the generalization contribution of the fast process quickly tunes toward the second force field, while the slow state still generalizes for the adaption of the first force field. During the spontaneous recovery phase, the generalization effect of the fast process quickly disappears, and the generalization, centered on the target location, washes out with the time constant of the slow process.

**Figure 1:**
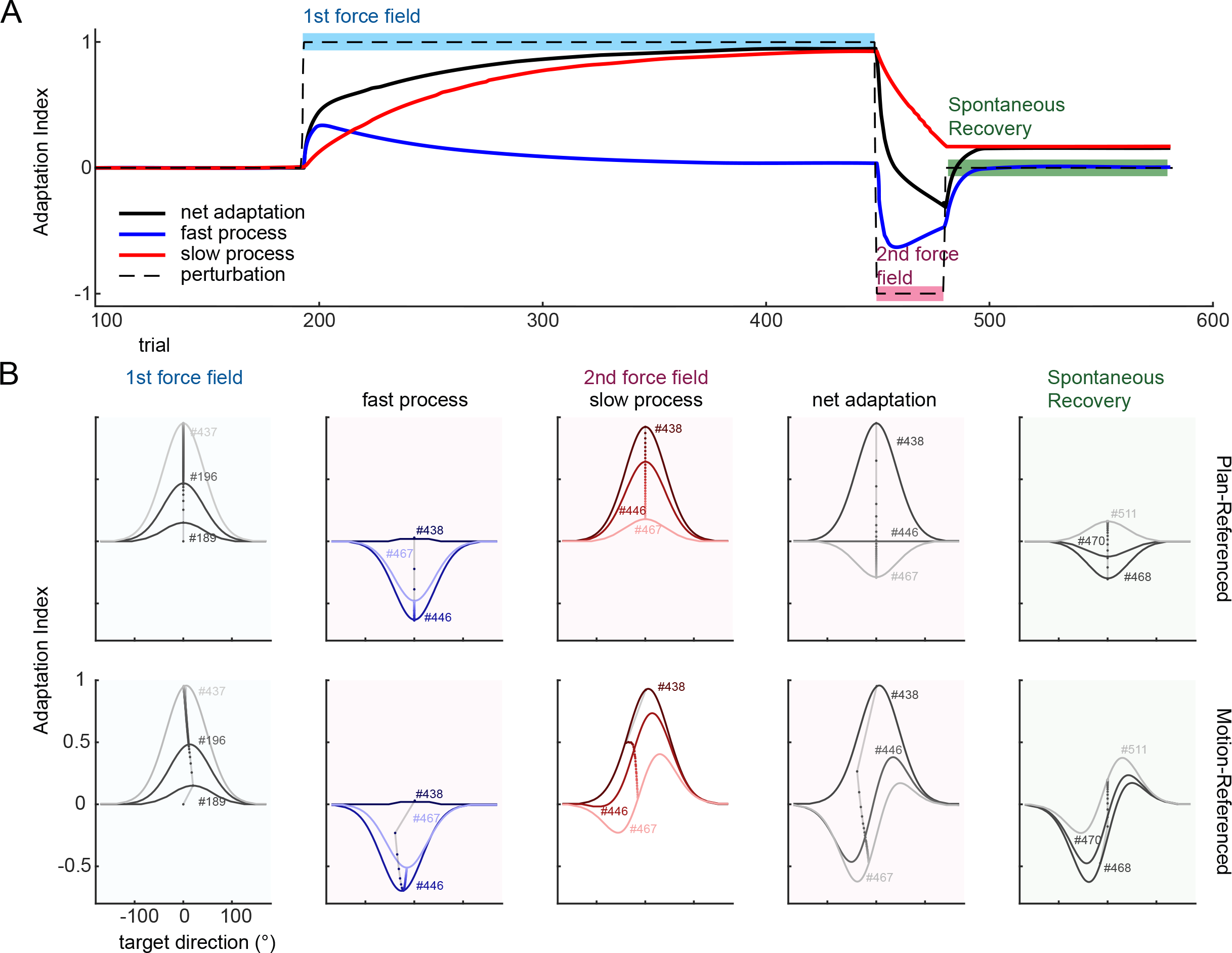
A: Spontaneous recovery paradigm. A long exposure to the first perturbation (turquoise background) is followed by a short exposure to a second, opposite, perturbation (magenta background). Subsequently, a set of Error-clamp trials (green background) is introduced to measure spontaneous recovery. The net adaptation (black solid line) is the sum of a fast process (blue solid line) and a slow process (red solid line). **B: Generalization patterns**. Predictions on the emergence and change of the generalization curves of the plan-referenced model (top) and motion-referenced model (bottom) during the first perturbation (turquoise background), second perturbation (magenta background), and Error-clamp phase (green background). The numbers next to the generalization curves indicate the associated trial number; the small dots represent the angle at which the update took place from trial to trial. The contributions of the fast and slow process to the net generalization curves are only shown during the second perturbation block, where the differences between the two models are most profound.

Instead, if the fast and the slow process both credit the actual motion, a different pattern of generalization emerges. The update of the motor primitives’ weights is centered on the actual movement, for both processes. As a result, with the reduction of target error, the overall generalization curve shifts in the direction of the target location. Because the slow and fast process have different dynamics, this leads to distinct generalization contributions, as seen in the bottom row of Fig 1B. As a result, during the spontaneous recovery phase the memory of the second force field of the fast process quickly decays, and reveals the asymmetrical generalization of the slow process. The present study examined these two model-based predictions.

## Methods

### Participants

Twenty-four right-handed participants gave written informed consent to take part in the study and received a financial compensation of 15€. One male participant was excluded, showing inconsistent behavior in the spontaneous recovery phase as compared to other participants (fully compensating for the second force field for the full phase); the final sample consisted of 23 individuals (15 female, 8 male), aged 18 to 34 years (m = 24.5). Participants had normal or corrected-to normal vision and had no known neurological or motor deficits. Right handedness was confirmed with the Edinburgh handedness questionnaire (Oldfield, 1971). The study was approved by the ethics committee of the Faculty of Social Sciences at Radboud University Nijmegen. Participants were randomly divided into two groups (a CCW-CW and a CW-CCW group, respectively; see below). Participation took about 70 minutes (64 minutes – 75minutes, experimental task only).

### Apparatus and Setup

Participants sat in front of a planar robotic manipulandum (Howard et al., 2009) that could generate forces at the handle. Seat belts across the shoulders restrained trunk movements. While holding the handle of the manipulandum, participants made reaches in the horizontal plane with their right hand. Their reaching arm was supported by an air-sled on a table, allowing near friction-less movements. Visual stimuli were presented in the plane of movement via a semi-silvered mirror, reflecting the display of an LCD screen. The mirror ensured that participants could not see their arm. A force transducer (F/T Nano, ATI Industrial Automation, Inc.) measured the force at the handle. Handle positions and forces were recorded at 1000 Hz.

Visual stimuli were presented against a black background and included the starting position (grey circle, radius 1 cm) and target position (yellow circle, radius 1 cm) of the movement. Hand position, derived from the handle position, was presented as a circular cursor (red circle, radius of 0.5 cm). The start position was at the center of the screen, ~30 cm away from the participants’ body. Eleven targets were presented on an imaginary semi-circle with a radius of 12 cm around the starting location. Target directions were at 0°, ±5°, ±10°, ±15°, ±35° and ±60°, with 0° representing the forward direction from the starting location, negative target directions representing counterclockwise directions, and positive directions representing positive target directions (Fig 2C). The target at 0° was also the training target (see below).

**Figure 2:**
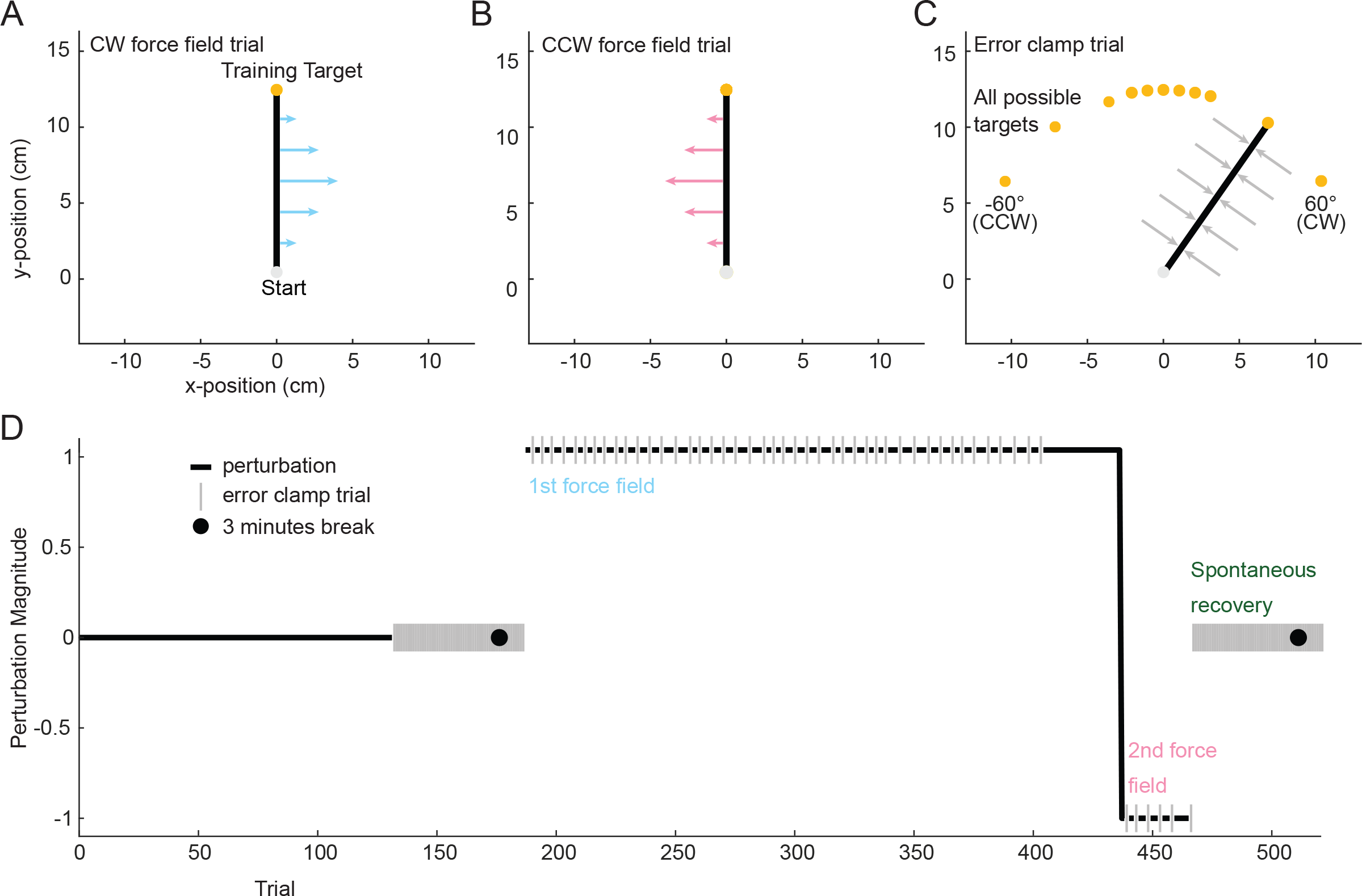
Experimental task and paradigm. **A, B**: **Clockwise (CW) and Counter Clockwise (CCW) force field trials** for the trained target direction. **C: Error-clamp trials** to quantify active compensation in terms of an Adaptation Index (AI) in the trained and untrained target directions. **D: Structure of a spontaneous recovery block** (repeated five times in the experiment) preceded by the familiarization phase, baseline Error-clamp 1 phase, break and Error-clamp 2 phase. Both in the 1^st^ force field and 2^nd^ force field phase, field exposure trials are interspersed with Error-clamp trials.

### Reach task

Before a trial started, participants were asked to place the hand cursor within the starting position, which then turned white. If the hand cursor remained within the starting position with a speed below 2.5 cm/s for 100ms, the target appeared. If participants did not start their movement within 600 ms after target appearance, a “React Faster” error message was displayed and the trial was repeated. If participants moved before target appearance, the trial was also aborted and repeated. Participants had to move through the target in one smooth movement, which exploded into multiple colored dots (‘visual fireworks’), accompanied with the word “correct” and a pleasant sound. If participants missed the target, the yellow target circle changed radius to include the hand position (radius equal to the distance of the hand to the target), which then slowly shrunk to the actual target size. This was accompanied by the word “accuracy”. Participants were encouraged to make smooth movements through the target with a duration in the range between 225 and 375 ms. If movement speed fell below 2.5 cm/s before the target distance had been covered, the trial was aborted, an unpleasant sound was played and the word “smoothness” was displayed. Movement duration was encouraged by providing target color feedback (red/blue), auditory feedback (same sound), and a displayed message (“Move slower”/”Move faster”).

The paradigm contained three types of reach trials: the robotic manipulandum could be off (*null-trial*) or produced a velocity-dependent curl force field (*force-field trial*). In the latter, force *F* depends on the hand velocity 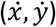, such that the force is perpendicular to the current movement direction and proportional to hand speed:

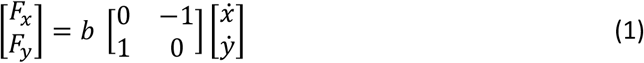

The damping constant (*b*) was set to 13 Ns/m and −13 Ns/m for a clockwise or a counterclockwise force field, respectively. The manipulandum could also produce an error-clamp (*Error-clamp trial*) in which the movement is constrained onto a straight line from the start position of the movement (hand speed > 2.5 cm/s) to the target position, to measure the compensatory force generated by the participant (Scheidt et al. 2000). The error-clamp trials used a stiffness of 3000 N/m and a damping constant of 25 Ns/m, and were used to determine the level of adaptation.

### Experimental paradigm

The experimental paradigm began with a familiarization phase of 132 null-trials, i.e., 12 reaches to each of the 11 target directions (Fig. 2D). This was followed by 44 Error-clamp trials (4 trials in each target direction) to establish a baseline adaptation index (see Analysis). After a 3 minutes break, participants completed the following sequence of experimental phases five times: Error-clamp 1 phase (11 trials, one for each pseudo-randomly selected target direction to check for savings), first force field phase (250 trials), second force field phase (15-50 trials), and Error-clamp 2 phase (44 trials, four for each target direction), and a 3 minutes break. Following Forano et al. (2020), the number of trials in the second force field phase varied among participants. This was done to make the Error-clamp 2 phase (the spontaneous recovery phase) start at an adaption level that showed no compensation to either force field. The sequence was repeated 5 times so that each target direction was tested once in the initial 3 trials of the spontaneous recovery phase, allowing to assess the generalization of the fast learning process. The fifth sequence did not end with a break, but was immediately followed by the Error-clamp 1 phase and 44 washout null-trials.

During the first force field phase (CW or CCW), participants performed 206 force field trials toward the 0° target, intervened by an Error-clamp trial after every four to six trials. The first five of these Error-clamp trials probed the targets close to the training direction (0, ± 5, ±10) to trace initial adaptation. The remaining 39 Error-clamp trials were interspersed pseudo-randomly across the full range of target directions to estimate generalization.

The second force field phase (opposite to the first force field) was organized in mini-blocks of 5 trials, containing 4 force field trials and one Error-clamp trial (randomly selected, but not the first trial) to the 0° target. The active compensatory force generated by the participant in these Error-clamp trials was quantified online (Forano & Franklin, 2020). If mean compensation over three mini-blocks changed sign (i.e., the initial compensation for the first force field had vanished and participants started to compensate for the second force field), the second force field phase was aborted, so yielding a different number of trials in the second force field across participants.

During Error-clamp 2 phase (the spontaneous recovery phase), the target directions were selected such that across the 5 sequence repetitions each target was at least once among the first three trials. For the remaining trials, target directions were selected pseudo-randomly such that each target direction had 4 repetitions in every sequence repetition.

### Analysis

Analyses were performed offline in MATLAB (2019b), based on raw data from the manipulandum. Start and end point of each reach were determined by taking the first and last timepoint at which 15% of the maximum speed of the forward movement was reached. For 17 trials (4 Error-clamp trials) this criterion could not determine a clear start and endpoint of the reach and start and end point were determined manually, or the trial was excluded (one Error-clamp trial).

To evaluate learning we used two measures: the angular kinematic error on the force field trials and the active force compensation in the Error-clamp trials for the 0° training direction. Angular error was taken as the direction of the hand at maximum speed relative to the forward direction, both referenced to the point where the hand left the starting position. For the active compensation in the Error-clamp trials we calculated the Adaptation Index (AI, Smith et al., 2006) by regressing, without an offset term and per trial, the perpendicular forces measured on the robot handle to the expected forces based on hand velocity in the channel (see Eq 1). Similarly, generalization was quantified in terms of active compensation in the Error-clamp trials for the non-trained target directions. Individual participant’s AI values were baseline corrected before being used in the model fitting procedure. Using the clamp trials from the baseline block (introduced after familiarization, 4 to each target), the average AI was computed per target. These averages were subtracted from the AIs in all other clamp trials tested for the same target.

### Model

Following the dual-rate model from Smith et al. (2006), learning occurs based on the difference between the actually experienced perturbation (*f*) in a trial (*t*) and the expected perturbation (*x*), resulting in an error (*ε*),

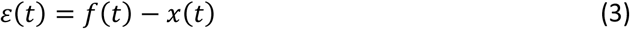

In the present experiment, *f* represents the experienced force, which was normalized to −1 for CCW perturbations and to 1 for CW perturbations.

The expected perturbation depends on the state of the fast and slow process at the start of a trial:

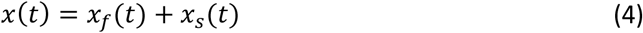

In the original model by Smith et al. (2006) the update of the fast and slow state are based on the retention (*r_n_*) of the previous state estimate and the learning (*l_n_*) from the error on the current trial *ε*(*t*):

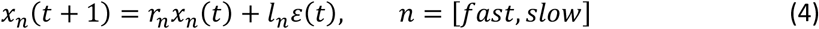

To simulate how the two states generalize across different movement directions, we reformulated this model as a combination of weighted motor primitives, each specified as a Gaussian-shaped tuning function (c.f. Yokoi et al., 2014). Hence, instead of directly updating state estimates, the weight vectors associated with the slow and fast state are updated, so that each determines the contribution of a set of motor primitives to the slow and fast state. The motor primitives are defined as a set (G) of 100 Von Mises distributions (*g)*, to ensure around the circle generalization:

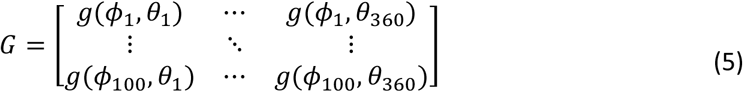

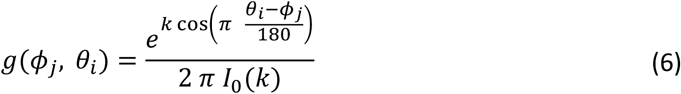

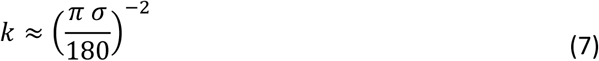

in which *ϕ_j_* is the preferred direction of a given motor primitive. The motor primitives were uniformly distributed from 0 − 2*π*. *θ_i_* is the movement direction, in degrees (discretized at one degree). I_0_(k) is the zeroth order modified Bessel function of *k*. The value of *k* was chosen such that the Von Mises approximated a Gauss curve with a width parameter σ of about 30°. These motor primitives represent the underlying neural tuning curves that convert a required movement direction into a required force. The weights associated with these primitives were updated separately for the slow and fast process:

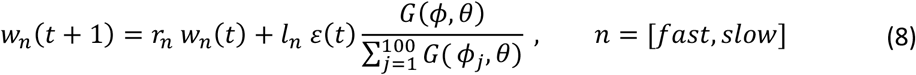

in which *θ* is the angle relative to which the weights are updated. This angle was taken as the planned movement direction (or target direction), (*θ* = *θ_target_*) for plan-referenced updating, or the actual movement direction of the hand (*θ* = *θ_hand_*) for motion-referenced updating. In Error-clamp trials we assume that no learning occurs since no error is experienced. Therefore, the update for the next trial only depends on the retention:

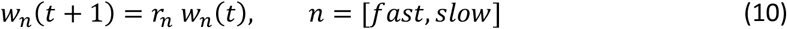

For theoretical and experimental reasons, the simulations above consider both processes to be either plan- or motion-referenced. We ignored the alternative combinations (plan-referenced for the one, and motion-referenced for the other, or vice versa) because their predictions only slightly deviate from the main models. These deviations are only apparent during the second perturbation, a phase in which we experimentally cannot probe generalization.

### Model fitting

The plan-referenced and motion-referenced model were fit to the baseline corrected AIs derived from the individual participant’s Error-clamp trials during the learning and spontaneous recovery phases in the experiment. Error-clamp trials of the ± 60° target directions were excluded due to unexpectedly high AI values. We used Bayesian Adaptive Direct Search (BADS; Acerbi & Ma, 2017) to minimize the sum of squared error between the model predictions and the AIs. The fast learning rate was constrained between 0.0001 < *l_fast_* < 0.9999 and the slow learning rate between 0.0001 < *l_slow_* < 0.15. The bounds of the slow learning rates were chosen strictly but realistically based on recovery analyzes in which prior simulations showed that a larger range led to multiple fit solutions. For the retention rates of the fast and the slow process, the bounds were 0.5 < *r_fast_*,*_slow_* < 0.9999. Next to these hard bounds, BADS allows for softer bounds on the most probable values of the parameters which were set to: 0.1 < *l_fast_* < 0.6, 0.5 < *r_fast_* < 0.8, : 0.0001 < *l_slow_* < 0.07, 0.95 < *r_slow_* < 0.9999. Starting values were randomly chosen within the probable bounds, and fits were repeated 10 times.

To validate our fitting procedure and set parameter constraints, we performed a recovery analysis. We generated 200 datasets with parameters within the probable bounds and added normally distributed noise (m = 0, SD = 0.1) to the observations. In the simulated and recovered data we implemented the updating angle by mapping the model error to an update angle relative to the target. For each parameter, the correlation coefficient between the simulated and recovered value was always >0.82. A recovery analysis with the participants’ parameter values as bounds for the generative model resulted in even higher correlations, > 0.93, suggesting that recovery is very good.

For comparison between plan- and motion-referenced updating models, we calculated BIC scores by using the mean squared error (MSE) per model over the Error-clamp trials during the learning and spontaneous recovery trials for each participant (Berniker et al., 2014). A group level BIC was taken by summing the BIC of all participants. We interpreted the difference in BIC following Kass & Raftery (1995): BIC > 10 very strong evidence, BIC >6 strong evidence, BIC >2 positive evidence.

#### Simulations of the fitted parameters in a spontaneous recovery paradigm

By using the best fitting parameters for both the plan- and motion-referenced model, we simulated the model predictions for individual participants during one sequence of the present paradigm (i.e. 132 familiarization trials, 44 baseline trials, 250 FF trials (CW), 15 FF trials (CCW), 44 Error-clamp trials). Note that although the number of trials in the second force field was variable across participants in the experiment, this simulation only used the initial 15 trials of the second force field. The simulations of the motion-referenced model used the angular error of the individual participants during the first block in the updating process. The mean model prediction for all possible target directions was calculated by taking the mean across the simulation of individual participants.

#### Model predictions vs data in the experimental design

To evaluate the predictions of the two models to the individual subject’s data, we simulated each model’s behavior with the best fitting parameters but now according to the subject specific trial structure encountered during their experimental task. To visually compare the model predictions and the data, we flipped the data and model predictions of the CCW-CW group (over the x-axis to match force compensations to the first and second force field, and over the y-axis to match update angles). We calculated the mean AI over blocks, participants (and trials) for relevant phases of the experiment (i.e. final part of the first and second force field and initial and final part of spontaneous recovery).

## Results

Participants performed 5 successive blocks of a spontaneous recovery paradigm. Apart from measuring learning in Error-clamp trials in the trained direction we also assessed generalization using Error-clamp trials in non-trained directions.

### Behavior

#### Adaptation dynamics

As shown in Figure 3A and B, during the first encounter with the first force field (block 1, blue dots), the participants’ initial reaches show large deviations from a straight path, resulting in large angular errors compared to baseline. These errors are in opposite directions for the CW-CCW (Fig. 3A, blue dots) and CCW-CW (Fig. 3B) participant groups. Over repeated trials, these errors diminish – a marker of adaptation. When the opposite force field is introduced (fuchsia dots), errors are initially particularly large, and of opposite sign, because participants are still actively compensating for the first force field. However, participants rapidly start adapting to this second force field. This error pattern repeats itself in later blocks (2-5), although the initial error in the first force field is at a much lower level.

**Figure 3:**
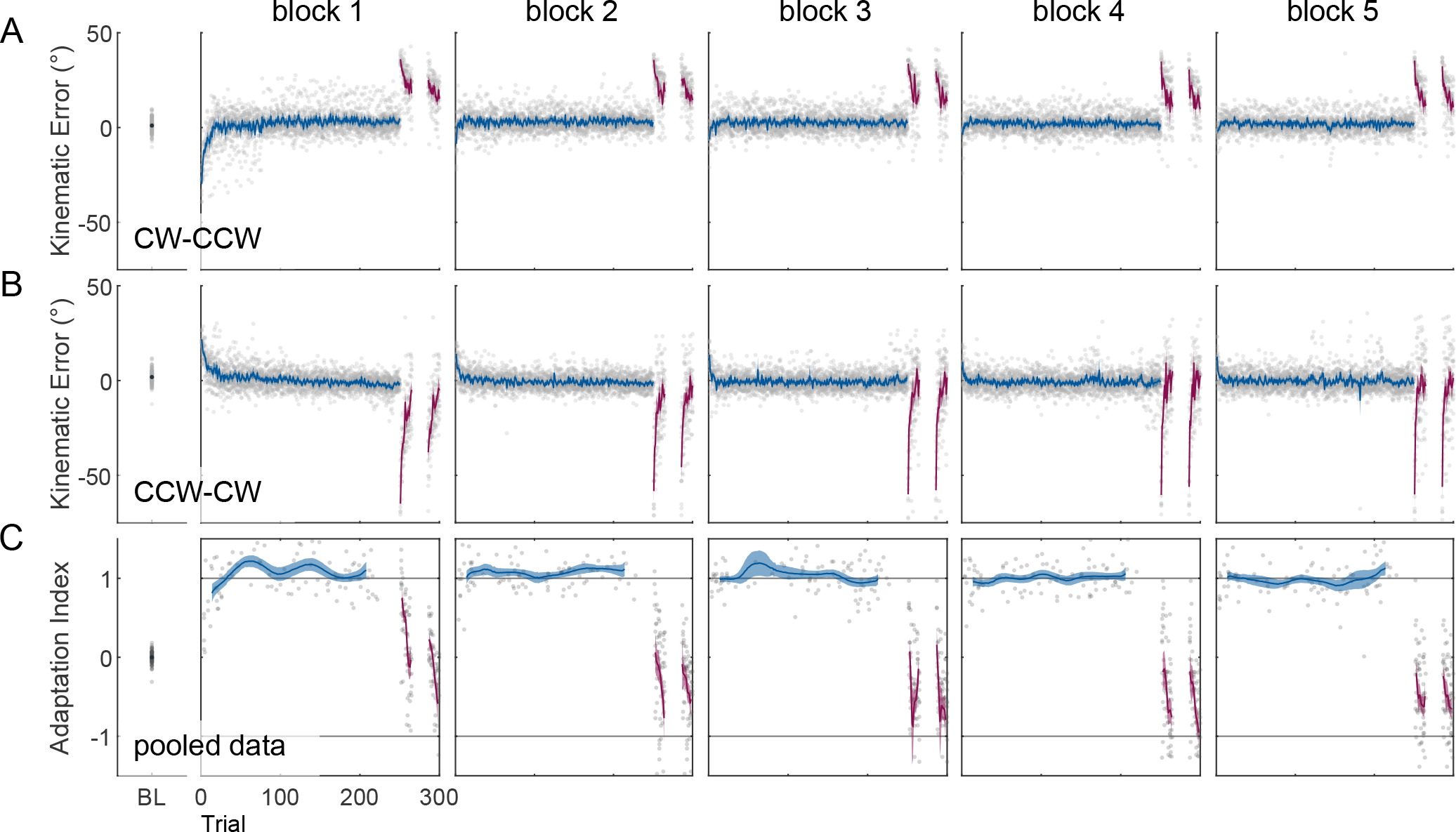
Force field adaptation for the trained target direction. **A, B: Kinematic error** (angular error at maximum movement speed) during the baseline (black dot) and force field exposures (block 1-5). The exposures to the first force field (blue lines) and second force field (pink lines) are depicted as mean ± SE. Data falling within 15 trials of the beginning (or end) of the second force field was aligned to first (or last) trial to indicate the initial (and final) level of kinematic error. Individual data points are depicted as light gray dots (0% and 1.1% of the data points of A and B fall out of the axis limits; A: N = 12, 7 female, B: N = 11, 8 female). **C: Active compensation in Error-clamp trials**. Baseline corrected adaptation index for the training target. The mean baseline (black dot), and sliding window mean (Hamming window) ± SE of the Error-clamp trials of the first force field (blue line, window size of 55 trials) and second force field (pink line, window size of 5 trials). Grey dots represent individual data points (4.3% of data points fall out of the axis limits). At the beginning and end of the curve the window size shrank on both sides to the amount of trials prior to the center of the sliding window. As in **A** and **B**, error clamp data falling within 15 trials from the first and final trial of the second force field were aligned either start or end to indicate initial and final AI levels.

Figure 3C corroborates these kinematic observations in terms of the adaptation index (AI). The AI is a measure of the compensatory force in the Error-clamp trials. During the baseline phase (BL), participants do not generate a compensatory force, as indicated by the mean AI (over participants and trials) around zero. For the subsequent force field blocks, each point represents a single Error-clamp trial of a single participant for the trained target direction. With the introduction of the first force field the AI rises rapidly to almost full compensation (blue dots). With the introduction of the second force field (fuchsia dots), the AI rapidly drops back to zero. The exposure to this second force field was abandoned as soon as the AI had dropped toward zero and thus depended on the participant’s behavior (see Methods). For most participants the spontaneous recovery phase started when the AI had just switched sign, but some had already started compensating for the second force field by the time the spontaneous recovery phase had started. In line with the results based on angular error, the pattern of the AI repeats itself in the subsequent blocks (2-5). The adaptation index increases more rapidly in the later blocks, which is in line with the reduction of initial error in the force field trials.

#### Generalization of the first force field

Figure 4 depicts how the active compensation for the first force field during each block generalizes to other untrained, movement directions, as assessed in intervening Error-clamps. For each target direction on the abscissa, the data of subsequent blocks is staggered horizontally (individual participant data in gray, participants’ mean in blue). The level of active compensation (AI) is more substantial and robust for movement directions close to the training direction (0°, ±5°) than further away (± 35°, ±60°). As the block number increases, the generalization pattern largely remains, although for some larger target angles (CCW-CW 35° and 60°), there seems to be a systematic reduction, and possibly a sign shift, of the adaptation index.

**Figure 4:**
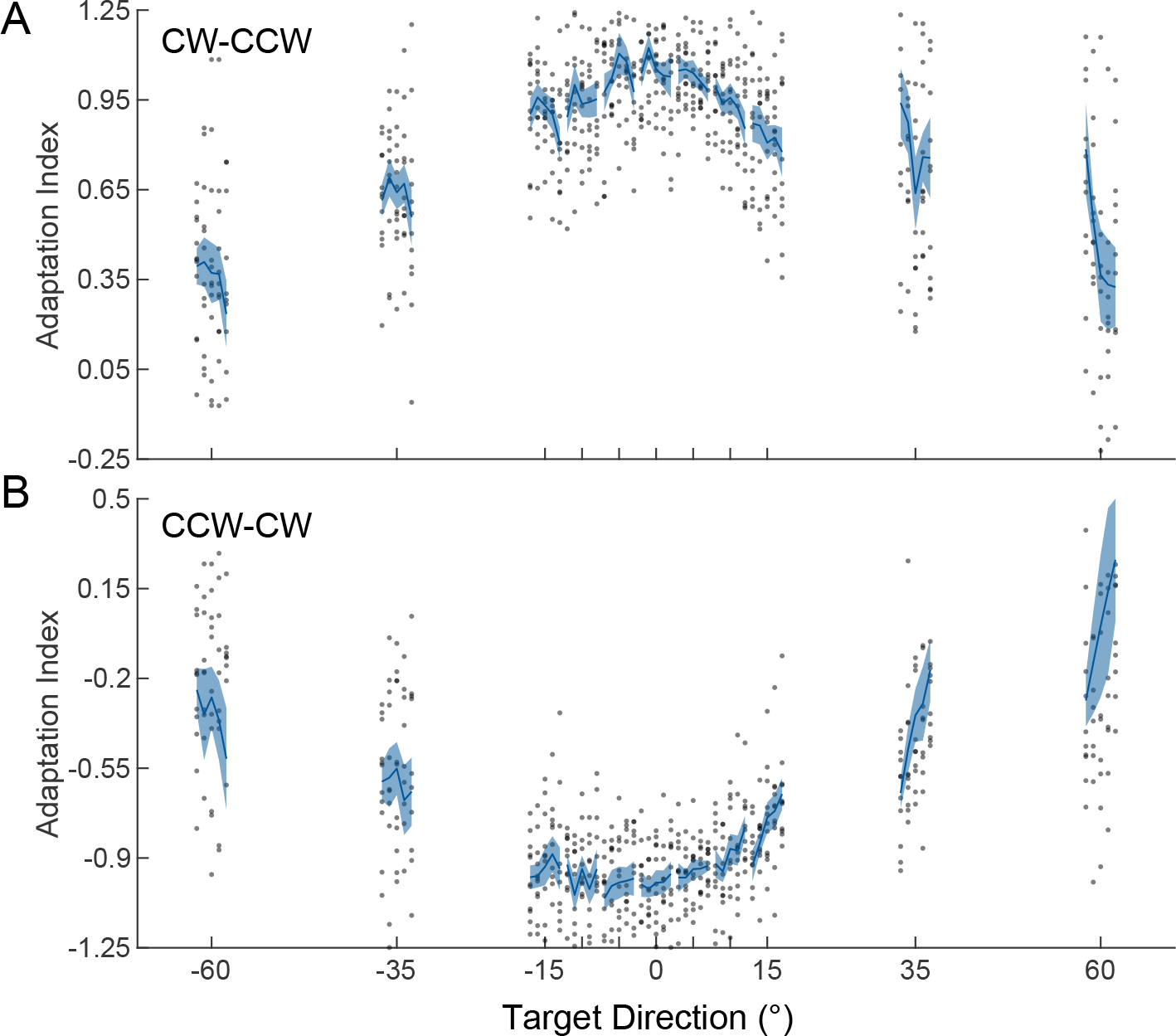
**Generalization of adaptation**, quantified in Error-clamp trials. Adaptation index (Mean ± *SE*) across participants and trials for the first force field. For each movement direction the lines represent, from left to right, the AI evolvement over blocks. Individual participant data (averaged over trials) are shown as grey dots (5.30% and 5.8% of the data points of A and B fall out of the plotting range; A: N = 12, 7 female B: N = 11, 8 female).

#### Generalization during spontaneous recovery

Figure 5 depicts the changes of adaptation index for trained and untrained directions during the spontaneous recovery phase for the two perturbation groups separately (upper and lower row, respectively) but pooled across blocks. Because movement directions are probed at different trials relative to the start of each spontaneous recovery phase, the mean is calculated over blocks, sets of 3 trials (relative to the start of the spontaneous recovery phase), and participants (green right-pointing triangle, number of trials ranging between 8 and 26). Across participants, the mean AI during spontaneous recovery demonstrates a clear rebound to compensating for the first force field, for both the trained direction and the untrained directions. Also note that the initial compensation as well as the strongest level of rebound are not centered on the trained target direction and are slightly asymmetrical between the two groups. To understand these experimentally observed patterns, we resort to fitting a plan-referenced and motion-referenced version of the dual rate learning and generalization model to the individual subject’s data, as described in the Methods.

**Figure 5:**
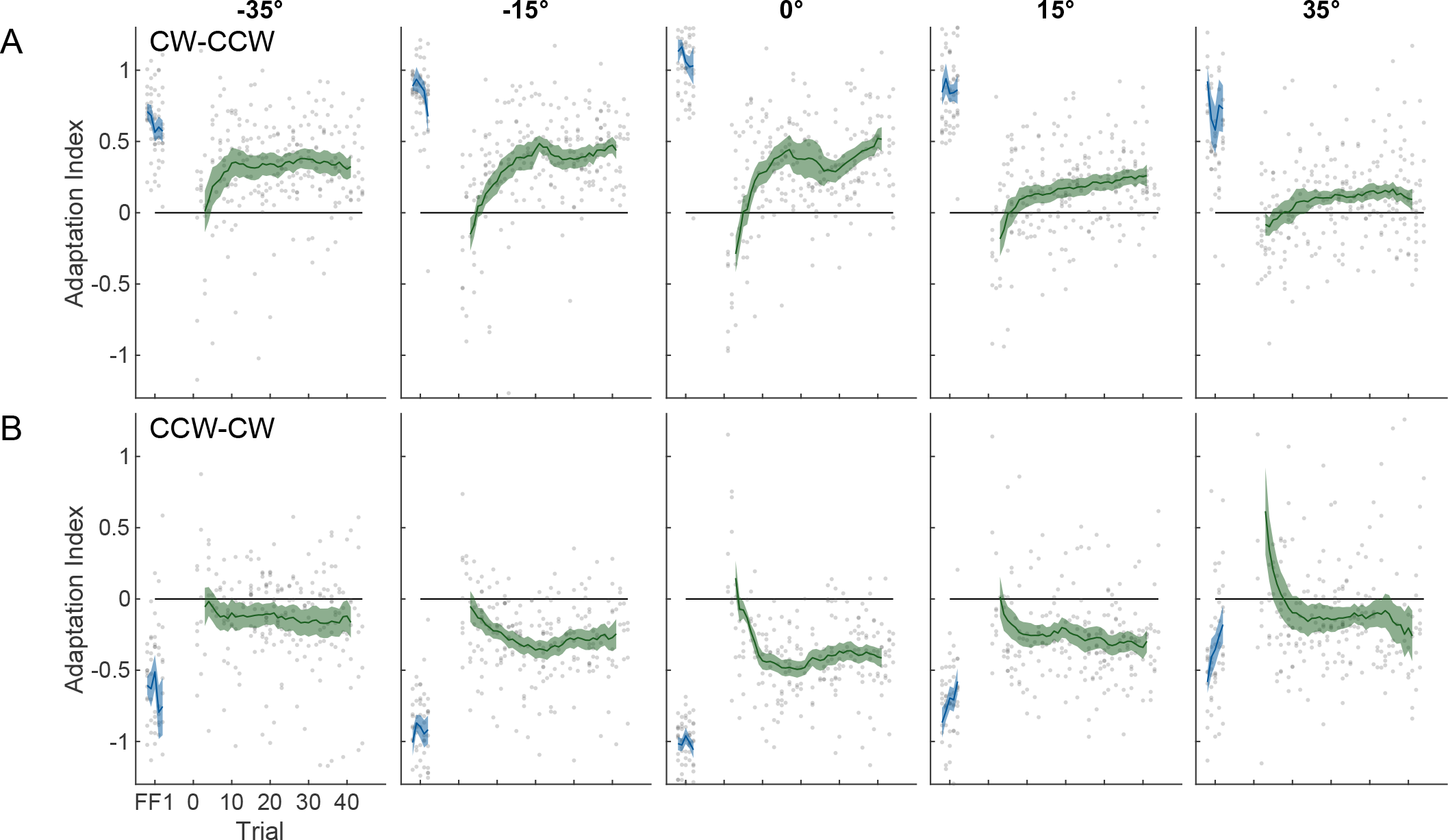
Force expression in the spontaneous recovery phase (Error-clamp 2 phase). Sliding window mean ±SE (in green, window size of 11 trials) compared to the mean ±SE AI at the end of the first force field (in blue) in the CW-CCW group (**A**: N = 12, 7 female) and CCW-CW group (**B**: N = 11, 8 female). Gray dots indicate individual participant data (2.2% and 1.7% of the data points of A and B fall out of the plotting range).

### Modelling

#### Model Fits

In both the plan- and motion-referenced model, force expression is formulated as a weighted readout of motor primitives and adaptation is achieved by updating these weights. The fast and slow adaptive process have their own weight vectors, which are updated independently. In the plan-referenced model, weights are updated relative to the planned motion (as defined by target direction), whereas in the motion-referenced model, weights are updated relative to a measure of the actual motion. Because of large variability of the data at the largest target angles (± 60°), we excluded these Error-clamp trials from the fits. We evaluated the performance of the two models based on a BIC analysis at the individual and group level.

Figure 6A shows the difference in BICs for the individual participants and at the group level. While at the group level the plan-referenced model best explained the data, at the level of the individual participant there were clear idiosyncratic preferences. More specifically, the BIC favored the plan-referenced model in 14 participants (one positively, two strongly, and eleven very strongly) and the motion-referenced model in 9 participants (three strongly, and six very strongly). Figure 6B-E shows the best-fit parameters for the two models for the individual participants. In terms of the best-fit parameters, the motion-referenced model yielded a higher learning rate of the fast process, the plan-referenced model a slightly higher learning rate of the slow process (see Figure 6). For the plan-referenced model, the retention rate of the fast process seems higher; the retention rate of the slow process appears slightly higher in the plan-referenced model.

**Figure 6:**
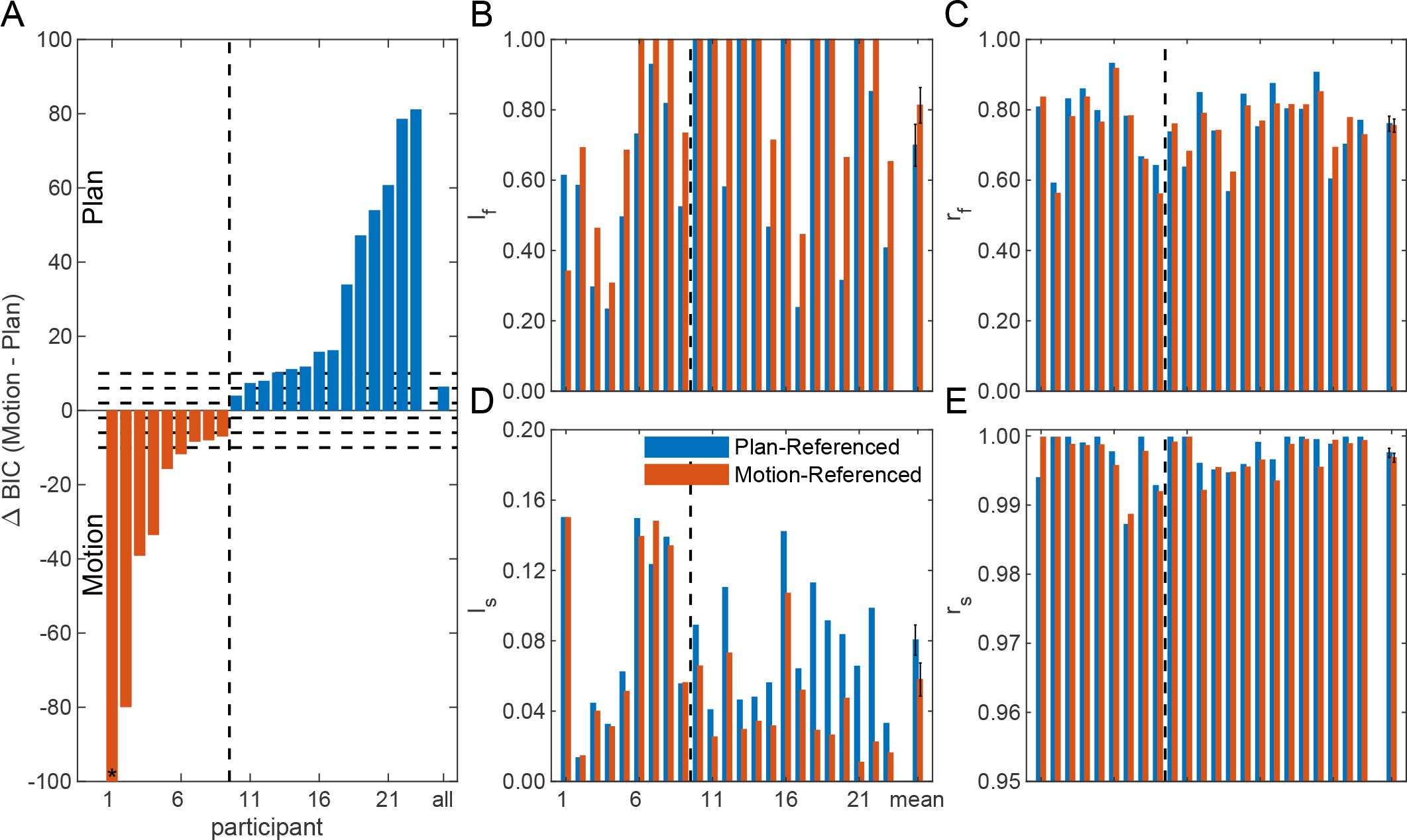
Comparison between plan- and motion-referenced updating model. **A: difference in BIC** (ordered by value). Orange bars indicate a difference in BIC favoring the motion-referenced model, blue bars indicate a difference in BIC favoring the plan-referenced model. *: bar reaches value of −230. **B-E**: show the parameter values of both model fits per participant ordered by the difference in BIC. **B**: learning rates of the fast process. **C**: learning rates of the slow process. **D**: retention rates of the fast process. **E**: retention rates of the slow process.

#### Plan- versus motion-referenced model predictions in a spontaneous recovery paradigm

To further scrutinize these differences in the model fits, we simulated the generalization of the slow and fast processes as well as the overall generalization of either model for the first block of the present paradigm, based on their best-fit learning and retention rates (across all participants). Figure 7A and B illustrate the generalization over movement directions as the adaptation evolves over trials. Warmer (red) colors indicate adaptation to the first force field, cooler (blue) colors reflect adaptation to the second force field. The plan-referenced model (Fig 7A) predicts a symmetric generalization profile around the central movement direction, with rapid initial adaption to the first force field mostly by the fast process, and an increasing contribution from the slow process over time. The white line indicates the direction relative to which the primitive weights are updated in this model. This is the central movement direction (i.e. the training direction of the adaptation) throughout the paradigm according to the plan-referenced model. The generalization curves of the fast and slow process, as well as the net adaptation at the time points of the dashed lines are depicted in Figure 7D (dashed lines). When the force field is reversed (second force field), the fast process quickly adapts (as shown by the cooler color), while the slow process largely retains its level of adaptation (also visible in Fig 7E, dashed lines). During the spontaneous recovery phase, the contribution of the fast process to the overall generalization diminishes, and the slow process dominates the observed generalization pattern, centered on the trained target direction (see also Fig 7F, dashed lines).

**Figure 7:**
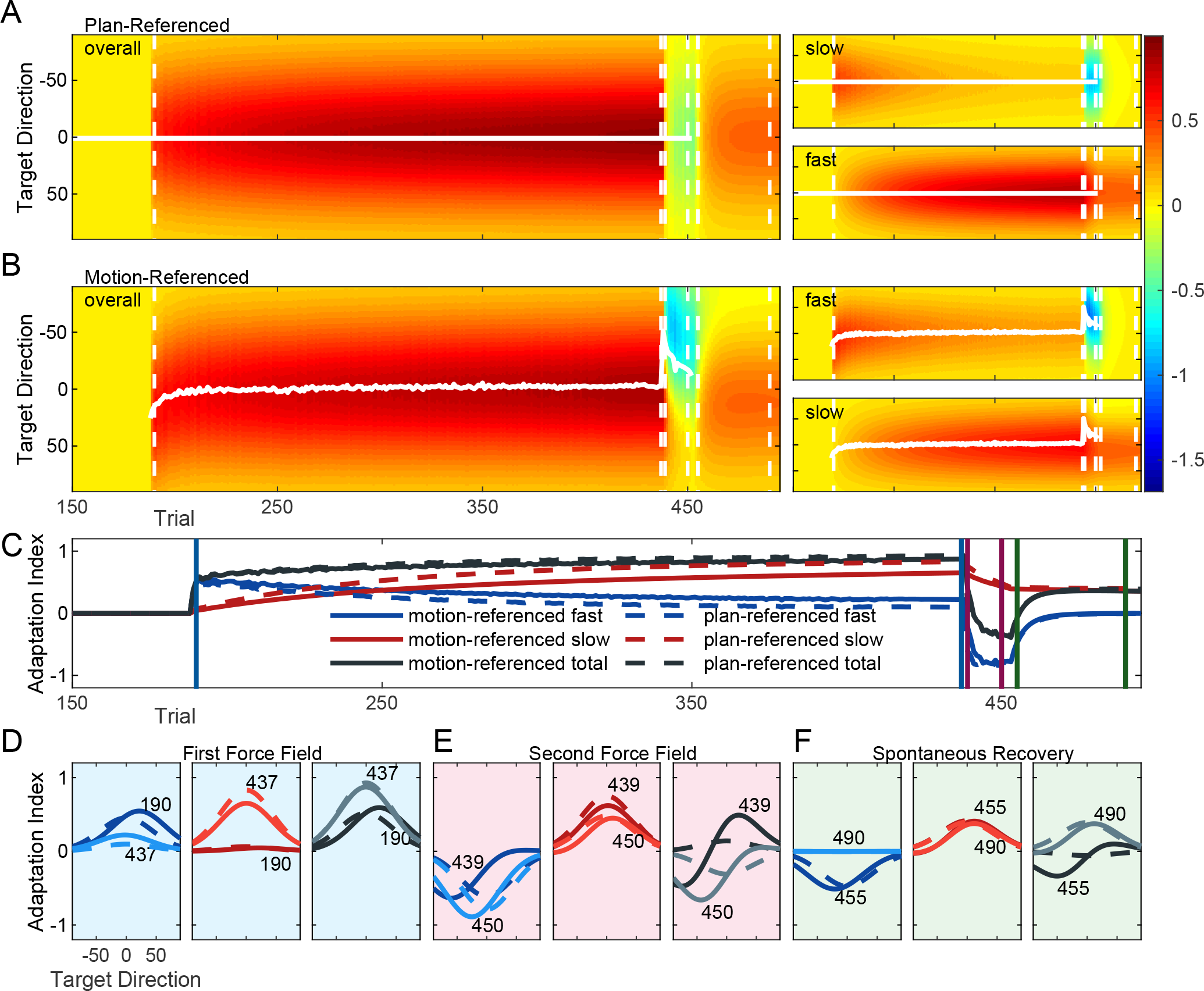
Mean model predictions in a spontaneous recovery paradigm. **A**, **B: Plan- and Motion-referenced model predictions over trials and target directions**: the left panels show the predictions of the total adaptation, the right four panels show the predictions of the fast and slow processes. White lines indicate the mean updating angle. Axis labels of all plots as in the overall plot of **B**. **C: Adaptation at training target direction**, predictions of the plan-referenced (dashed) and motion-referenced (solid) model, for the fast (blue), slow (red) process, and total adaptation (black). **D: Generalization over target directions at different learning phases**, generalization of early (stronger colors) and late (lighter colors) trials in each phase (1^st^ force field, 2^nd^ force field, and spontaneous recovery; see vertical lines in **C**) of the fast (blue), slow (red) process, and the total adaptation (black) are shown per model (motion-referenced in solid and plan-referenced in dashed lines) over target directions.

In contrast, the best-fit motion-referenced model predicts asymmetric generalization profiles, as shown in Fig 7B. Because the movement direction defines the update direction of the primitive weights, the generalization is initially skewed towards this direction but readily, with trials, shifts to the central target direction when movement and thus adaptation aligns with the planned movement direction. This shift is most prominent for the fast process. Again, panels D, E and F show (solid lines) the generalization curves at the different time points indicated by the dashed white lines in B. Because there is variance in the update direction of the motor primitives (because it is based on angular error), the predicted generalization curves for the fast and slow processes, as well as for the overall adaptation, are slightly wider than for the plan-referenced model. The slow process of the motion-referenced model also reaches a lower level of compensation for the first force field (as compared to the plan-referenced model, see Fig. 7C). To reach a similar level of net adaptation, the fast process shows higher levels of compensation for the first force field. This is due to the differences in learning and retention rates of the best model fits. During the second force field, the motion-referenced model diverges substantially from the plan-referenced model, as the largest update of the motor primitives occurs for those aligned with the actual motion, yielding rather asymmetric generalization profiles for both fast and slow processes, as well as the overall adaptation (see also Fig 7E, solid lines). At the level of the overall adaptation, these asymmetries result in a rapid decrease of generalization for negative target angles, whereas compensation for the original perturbation remains somewhat preserved for positive target angles. During the spontaneous recovery phase, the compensation for the second force field is quickly lost by the fast process, and the memory of the slow process becomes visible, which with trials starts to peak off-center because of the unlearning effects of the slow process of the second force field.

Note that these simulations do not suffer from the experimental impracticality that only one target direction can be tested per trial. Furthermore, the order of the targets in which the generalization was probed differed across participants. To compare model predictions to the individual participant’s experimental designs and data, we simulated both models in the participant specific experimental design, and compared their means over blocks, trials and participants.

#### Comparing Model Fits to the Experimental Data

Figure 8 presents the individual participants’ data and model predictions split by the winning model, i.e. the plan-referenced model for 14 participants and the motion-referenced model for 9 participants. The net generalization of the first force field (last Error-clamp trial for each target direction) is shown by the blue dots as the average values of the (simulated) AI for a specific target angle. The compensation of the last Error-clamp trial during the second force field is depicted in fuchsia. The trials of the spontaneous recovery phase, shown in green, are separated into the initial part (3 trials), in which the compensation for the second force field is most pronounced, and asymptote (last 22 trials, see Fig. 5), in which the memory of the compensation for the first force field, i.e. the spontaneous recovery, is most pronounced. Open symbols show data and simulations for the individual participants (average over trials and blocks), whereas filled symbols show the average across participants as well.

**Figure 8:**
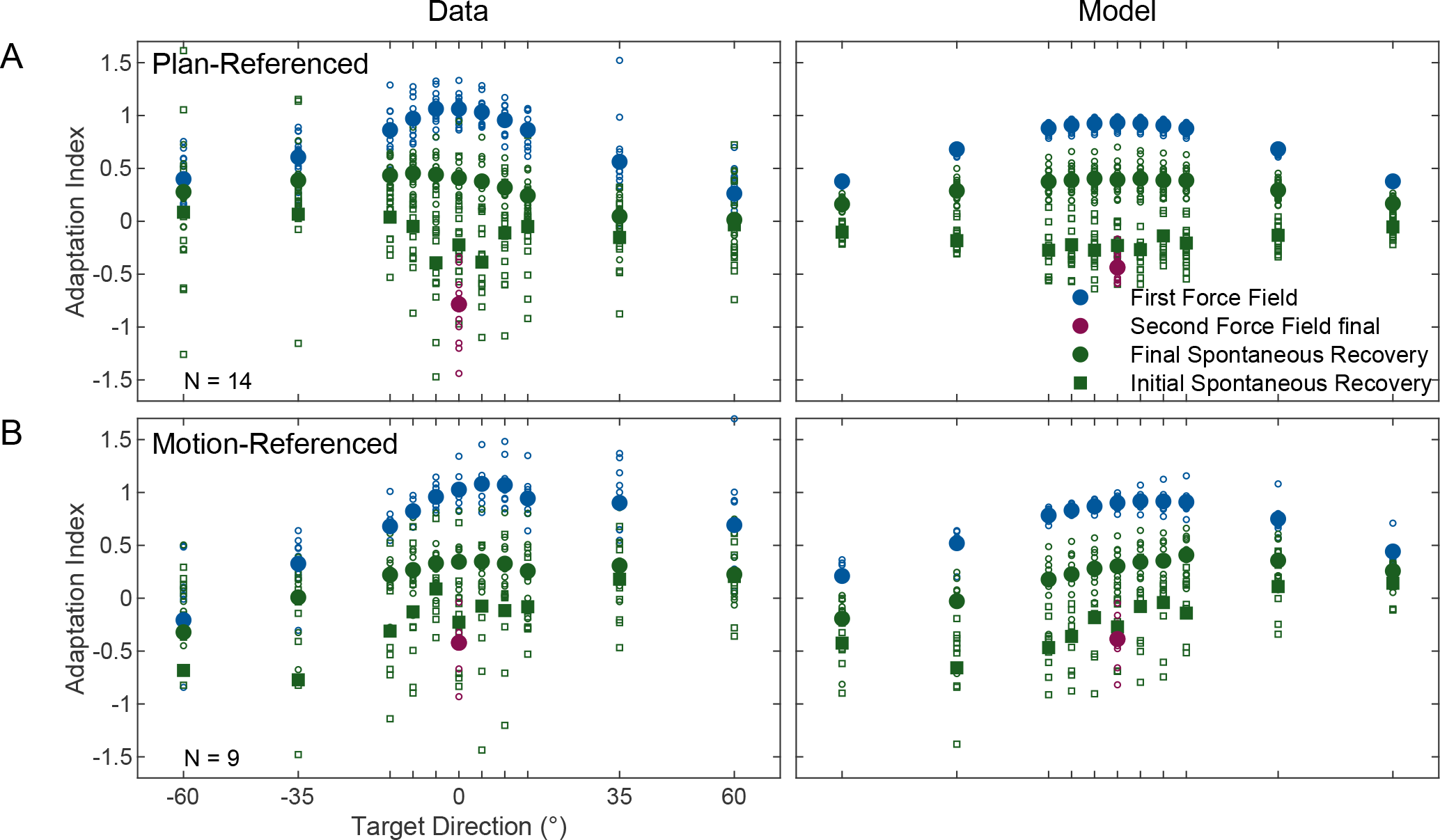
Data and model predictions split by the best explaining model. **A. Plan-referenced model**. Left panel, data from nine CW-CCW (two male, seven female) and five CCW-CW (two male, three female); right panel: model fits**. B: Motion-referenced model**. Left panel: data from three CW-CCW (all male) and six CCW-CW (one male, 5 female); right panel: model fits. Open symbols are the means over trials and blocks per movement angle and per participant. Filled symbols are the means over participants, blocks and trials per movement angle (1.6% and 0.2% of the data points of A and B fall out of the plotting range).

Figure 8A shows the experimentally derived AIs for the participants that showed generalization that best matched the plan-referenced model, whereas figure 8B shows the corresponding model predictions. The generalization for the initial force field (blue circles) is nearly symmetric in the data, as predicted by the plan-referenced updating model. When examining the patterns in the spontaneous recovery phase (green squares and circles), the participant data shows asymmetries, which are not captured by the plan-referenced model by design. These asymmetries are, however, opposite to what would be expected from the motion-referenced model (compare trends Fig 8A data, with Fig 8D model predictions). In particular, the rebound (green circles) toward the level of compensation in the first force field is lower at CW movement directions, the direction to which the hand was pushed during the first force field.

Figure 8C shows the data for the participants who showed generalization that best matched the motion-referenced model. These data show strong asymmetries in generalization during the first force field, with higher Ais in the movement directions corresponding with the actual hand trajectories. Also, the initial part of the spontaneous recovery phase (green squares) shows a clear asymmetry, with stronger adaptation (more negative AI) to the second force field for negative target angles, i.e. for directions that the hand was pushed toward by the force field, than for positive target angles. In addition, the rebound (green circles) toward the AI observed in the first force field is also strongest for the −35° direction. These asymmetries in the generalization patterns are in line with the predictions from the motion-referenced model (Fig. 8D).

## Discussion

It is generally accepted that motor adaptation is governed by multiple interactive underlying processes, operating on different timescales (Lee & Schweighofer, 2009; Smith et al., 2006), although there may be exceptions for some tasks (Ingram et al., 2011; Rudolph et al., 2020). Here, using a spontaneous recovery paradigm and model-based analyses, we tested how the adaptative states associated with these processes generalize to other movement directions over the course of learning to reach through a velocity-dependent force field. Following Yokoi et al. (2011, 2014), we extended the two-state adaptation model of Smith et al. (2006) with a set of motor primitives, whose readout depends on a weight vector that can be updated. Adaptation in this model is achieved by updating individual weights of the fast and slow adaptive process separately based on the observed error. Depending on whether this error-based updating is centered on the planned motion (Donchin et al., 2003) or the actual motion (Gonzalez Castro et al., 2011), the model predicts distinct contributions of the slow and fast adaptation process to the overall generalization function. Comparing the empirical data to the predictions of these two model variants, our participant population fell within a continuum of evidence for plan-referenced to evidence for motion-referenced updating.

For both models we used a fixed set of 100 motor primitives, uniformly distributed around a full circle. Their preferred directions, number, and tuning widths were based on earlier studies (Yokoi et al., 2011, 2014). We ignored the possibility that the fast and slow process could rely on motor primitives of different tuning widths, or that tuning properties could change over the course of adaptation (Cherian et al., 2013; Li et al., 2001; Padoa-Schioppa et al., 2004; Rokni et al., 2007), given that recent neurophysiological reports suggest that force field adaptation changes recruitment of neurons in motor cortex, not their tuning properties (Gallego et al., 2020; Perich et al., 2018; Perich & Miller, 2017). In the same vein, a recent study using population-based analysis examined the preparatory activity in motor cortex when monkeys learned to reach in an altered force environment (Sun et al., 2022). During adaptation, it was found that the population activity moved in dimensions orthogonal to those before the force environment was altered, with the same time course by which the monkey learnt to compensate for the altered force. Hence, this re-organization of the population dynamics could be a mechanism by which the brain learns and retains new motor behaviors.

To find signatures of the underlying adaptive processes to the overall generalization, our participants performed five successive blocks of a spontaneous recovery paradigm, demonstrating the re-expression of the initial adapted state, after it was followed by short reverse-adaptation in each block (Smith et al., 2006). Indeed, while there were single peaked generalization profiles in the Error-clamps during the first force field, during the spontaneous recovery phase the generalization function switched from compensation for the second force field to compensation for the first force field, for both trained and untrained movement directions. However, there was large inter-subject variability for motion- or plan-referenced updating, suggesting that participants may rely on different reference frames in the updating of their motor primitive weights. In the following paragraphs we will discuss possible explanations for these results.

In contrast to our mixed findings, participants in the study by Gonzalez Castro et al. (2011) showed motion-referenced updating at the group level. To examine possible differences introduced by the difference in experimental paradigm, we simulated their experiment 1 using our primitive-based motion-referenced updating model. In that experiment participants were exposed to a force field that switched between CW and CCW directions approximately every seven trials. After extensive exposure of our model to this switching pattern, the overall generalization curve indeed looks like their experimentally determined curve (see figure 2 in their report). The simulation further shows that this curve is largely driven by the slow process, because the fast process quickly washes out when the field direction changes.

In our paradigm, the long blocks of the first force field reveal a more robust generalization, centered at the target location, which is predicted by both the plan-referenced and the-motion-referenced model (see Figs. 4 and 8). The behavioral changes during the second force field are mostly driven by the fast process (Fig. 6) and the slow process may barely have changed because of the short duration of this perturbation. It cannot be excluded that this has obscured a motion-referenced generalization pattern in the spontaneous recovery phase in some participants. An alternative explanation for why we did not find a consistent reference frame across all participants could relate to the structure of the present task, which may have evoked different strategies among participants.

To expand on the latter, visuomotor adaptation studies with reaching movements have associated the fast process to an explicit strategy (Day et al., 2016; Heuer & Hegele, 2011; McDougle et al., 2015; Taylor et al., 2014). In these studies, participants indicated the aiming direction of their intended movement prior to executing it. The implicit contribution was then inferred as the difference between the actual reach direction and the explicit aim direction. Along with a two-state modeling analysis, it was shown that the fast process closely resembles explicit learning and the slow process approximates implicit learning (McDougle et al., 2015). Furthermore, the explicit component was found to generalize more globally across the workspace, while the implicit component generalizes more locally, around the explicitly aimed direction (Day et al., 2016; Heuer & Hegele, 2011; McDougle et al., 2017). This is in line with the modeling results from Tanaka et al. (2012) showing that the fast process is responsible for trial-by-trial generalization in visuomotor adaptation and the slow process is mainly responsible for the post-adaptation generalization. If these findings apply equally well to force field learning, it could be argued that in the task of Gonzalez Castro et al. (2011), the frequent switching of the force field direction probably pushed participants away from an explicit strategy and participants primarily learned implicitly based on the motion of their hand. In contrast, in the present paradigm there was no clear constraint for using an explicit or implicit strategy: some of our participants may have relied on an explicit aiming strategy, consciously pushing into the force field (Schween et al., 2020), whereas others may have relied predominantly on a motion-referenced implicit strategy. Future research could investigate how the explicit awareness of the task structure in different paradigms affects individual’s choices in the use of more explicit strategies.

As a final point of discussion, the participants in our study seemed to learn the compensation for the second force field more rapidly across the five spontaneous recovery blocks (Fig. 3). This may suggest that there is a third, even slower, adaptation process that retains information across blocks (Forano & Franklin, 2020). Alternatively, it has recently been suggested that a single motor memory only consists of a single, adaptable, state, but that at any moment in time a weighted average of motor memories is expressed and adapted based on the inferred context (Heald et al., 2021). This interpretation diverges significantly from the dominant notion of multiple adaptive states that override each other to form and express a single motor memory. Instead, contextual inference proposes that the brain builds separate memories for the null, CW and CCW force fields and over the blocks learns to infer the correct context more quickly. Although this framework has not yet been implemented as a network of motor primitives, it would be an interesting step for future work to test whether it can provide a unifying account for behavioral and neurophysiological observations in motor learning, including the generalization effects of the present study.

## Acknowledgements

This work was supported by an internal grant from the Donders Centre for Cognition. Support comes also from the research programme National Research Agenda, which is (partly) financed by the Dutch Research Council (project number 1292.19.298).

## Notes

### Competing Interest Statement

The authors have declared no competing interest.

### Summary of Updates

Data presentation has been substantially improved

